# Inbreeding parents should invest more resources in fewer offspring

**DOI:** 10.1101/062794

**Authors:** A. Bradley Duthie, Aline M. Lee, Jane M. Reid

**Affiliations:** Institute of Biological and Environmental Sciences, School of Biological Sciences, Zoology Building, Tillydrone Avenue, University of Aberdeen, Aberdeen AB24 2TZ, United Kingdom

**Keywords:** Inbreeding, inclusive fitness, mate choice, parental investment, relatedness, reproductive strategy

## Abstract

Inbreeding increases parent-offspring relatedness and commonly reduces offspring viability, shaping selection on reproductive interactions involving relatives and associated parental investment (PI). Nevertheless, theories predicting selection for inbreeding versus inbreeding avoidance and selection for optimal PI have only been considered separately, precluding prediction of optimal PI and associated reproductive strategy given inbreeding. We unify inbreeding and PI theory, demonstrating that optimal PI increases when a female's inbreeding decreases the viability of her offspring. Inbreeding females should therefore produce fewer offspring due to the fundamental trade-off between offspring number and PI. Accordingly, selection for inbreeding versus inbreeding avoidance changes when females can adjust PI with the degree that they inbreed. In contrast, optimal PI does not depend on whether a focal female is herself inbred. However, inbreeding causes optimal PI to increase given strict monogamy and associated biparental investment compared to female-only investment. Our model implies that understanding evolutionary dynamics of inbreeding strategy, inbreeding depression, and PI requires joint consideration of the expression of each in relation to the other. Overall, we demonstrate that existing PI and inbreeding theories represent special cases of a more general theory, implying that intrinsic links between inbreeding and PI affect evolution of behaviour and intra-familial conflict.

## Introduction

Inclusive fitness theory identifies how natural selection will act at any given level of biological organisation (Grafen, 2006). It thereby provides key evolutionary insights (Fisher et al., 2013; Bourke, 2014; Gardner and West, 2014; Liao et al., 2015), perhaps most iconically explaining self-sacrificial behaviour of focal individuals by accounting for the increased reproductive success of related beneficiaries that carry replica copies of their alleles (Hamilton, 1964*a*, *b*; Frank, 2013). Inclusive fitness theory also identifies relatedness as a central modulator of sexual conflict over mating and fertilisation (Parker, 2006), and of conflict among parents and offspring over parental investment (hereafter ‘PI’; Trivers, 1972, 1974; Kölliker et al., 2015).

Recently, there has been considerable theoretical and empirical interest in how relatedness affects sexual conflict (e.g., Pizzari and Gardner, 2012; Carazo et al., 2014; Chippindale et al., 2015; Faria et al., 2015; Martin and Long, 2015). In the context of sexual conflict over mating decisions, it remains somewhat under-appreciated that individuals can increase their inclusive fitness by inbreeding. Selection for inbreeding tolerance or preference is therefore sometimes predicted despite decreased viability of resulting inbred offspring (i.e.,“inbreeding depression”, hereafter ‘ID’; Parker, 1979, 2006). Furthermore, inclusive fitness theory pertaining to inbreeding versus inbreeding avoidance has focused solely on individuals' mating decisions, assuming no concurrent modulation of PI or offspring production (Parker, 2006; Kokko and Ots, 2006; Duthie and Reid, 2015). Such theory ignores that parents might be able to increase the viability of their inbred offspring through PI, potentially mitigating ID. Because parents are more closely related to inbred offspring than they are to outbred offspring (Trivers, 1974; Lynch and Walsh, 1998; Reid et al., 2016), inclusive fitness accrued from viable inbred offspring should be greater than that accrued from viable outbred offspring. Consequently, the inclusive fitness consequences of inbreeding and PI cannot be independent. However, to date, inbreeding and PI theory have been developed separately, potentially generating incomplete or misleading predictions concerning reproductive strategy.

A basic inclusive fitness model has been developed to predict female and male inbreeding strategies, wherein a focal parent encounters a focal relative and chooses to either inbreed or avoid inbreeding (Parker, 1979, 2006; Kokko and Ots, 2006; Duthie and Reid, 2015). If the focal parent inbreeds, then the viability of resulting offspring decreases (i.e., ID), but the offspring will inherit additional copies of the focal parent's alleles from the parent's related mate. The focal parent can thereby increase its inclusive fitness by inbreeding if the number of identical-by-descent allele copies in its inbred offspring exceeds the number in outbred offspring after accounting for ID (Parker, 1979, 2006; Kokko and Ots, 2006; Szulkin et al., 2013; Duthie and Reid, 2015). The magnitude of ID below which inbreeding rather than avoiding inbreeding increases a parent's inclusive fitness is sex-specific, assuming stereotypical sex roles. This is because female reproduction is limited by offspring produced, while male reproduction is limited only by mating opportunities. Hence, a focal female only indirectly increases the number of identical-by-descent allele copies inherited by her offspring when inbreeding, while a focal male directly increases the number of identical-by-descent allele copies inherited by siring additional inbred offspring. All else being equal, males but not females therefore benefit by inbreeding given strong ID, while both sexes benefit by inbreeding given weak ID (Parker, 1979, 2006; Kokko and Ots, 2006; Duthie and Reid, 2015). These predictions are sensitive to the assumption that there is a low or negligible opportunity cost of male mating. If inbreeding instead precludes a male from siring an additional outbred offspring, such as when there is an opportunity cost stemming from monogamy and associated biparental investment in offspring, then inbreeding is never beneficial (Waser et al., 1986). However, existing theory that considers these inclusive fitness consequences of inbreeding assumes that PI is fixed despite resulting variation in parent-offspring relatedness (Trivers, 1974; Lynch and Walsh, 1998; Reid et al., 2016).

Meanwhile, a separate general framework for PI theory, which typically (implicitly) assumes outbreeding, is well-established. Here, PI does not simply represent raw resources provided to an offspring, but is defined as anything that a parent does to increase its offspring's viability at the expense of its other actual or potential offspring (Trivers, 1972, 1974). One key assumption of PI theory is therefore that the degree to which a parent invests in each offspring is directly and inversely related to the number of offspring that it produces. A second key assumption is that offspring viability increases with increasing PI, but with diminishing returns on viability as more PI is provided. Given these two assumptions, the optimal PI for which parent fitness is maximised can be determined, as done to examine the magnitude and evolution of parent-offspring conflict over PI (Macnair and Parker, 1978; Parker and Macnair, 1978; Parker, 1985; De Jong et al., 2005; Kuijper and Johnstone, 2012). Such models assume that offspring are outbred, or have been specifically extended to consider self-fertilisation (De Jong et al., 2005). However, biparental inbreeding is commonplace in wild populations, directly affecting both offspring viability and parent-offspring relatedness (Trivers, 1974; Lynch and Walsh, 1998; O'Grady et al., 2006; Charlesworth and Willis, 2009; Reid et al., 2016). Such inbreeding might profoundly affect reproductive strategy if parents that inbreed can mitigate ID through increased PI.

We unify two well-established but currently separate theoretical frameworks; the first predicts thresholds of ID below which focal parents increase their fitness by inbreeding rather than by avoiding inbreeding (Parker, 1979), and the second predicts optimal PI given outbreeding (Macnair and Parker, 1978). By showing how inbreeding and PI decisions are inextricably linked with respect to their effects on inclusive fitness, we provide a general framework that identifies the direction of selection on reproductive strategy arising in any population of any sexual species. First, to demonstrate the key concepts, we focus on the reproductive strategy of an outbred, but potentially inbreeding, female that is the sole provider of PI. We show that her optimal PI changes predictably with her relatedness to the sire of her offspring, and with ID. Additionally, we model the consequences of a focal female being inbred for optimal PI and inclusive fitness. Second, we extend our framework to consider the consequences of strict monogamy, and associated obligate biparental investment, for optimal PI and inclusive fitness. Within both frameworks, we additionally show how inclusive fitness changes when focal females and monogamous pairs cannot adjust their PI optimally with inbreeding (e.g., if individuals cannot recognise kin, Penn and Frommen, 2010), as is implicitly assumed in all previous inbreeding theory (Parker, 1979, 2006; Waser et al., 1986; Kokko and Ots, 2006; Duthie and Reid, 2015).

## Unification of inbreeding and PI theory

We consider a focal diploid parent (hereafter assumed to be a stereotypical female) that can adjust the degree to which she invests in each offspring to maximise her own inclusive fitness, defined as the rate at which she increases the number of identical-by-descent allele copies inherited by her viable offspring (*γ*). This definition of fitness differs from previous models of PI (Macnair and Parker, 1978; Parker and Macnair, 1978), which instead define fitness as the rate at which viable offspring are produced and therefore cannot account for inclusive fitness differences between inbred and outbred offspring. We assume that offspring viability increases with increasing PI (*m*), with diminishing returns as *m* increases (following Parker and Macnair, 1978). Females have a total lifetime PI budget of *M*, and therefore produce *n* = *M*/*m* offspring, modelling the fundamental trade-off between the number of offspring produced and investment per offspring. We assume for simplicity that *M* » *m* (following Parker, 1985), but this assumption should not affect our general conclusions. Given these minimal assumptions, we can conceptually unify inbreeding and PI theory through a general framework that predicts the number of identical-by-descent copies of a female's alleles that are inherited per offspring (*ζ*_off_), scaled relative to the female's kinship to herself,

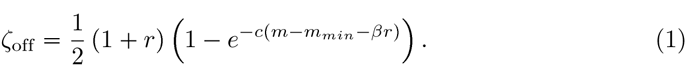

Our model in Eq. 1 can be understood in two pieces (parameters are summarised in Table 1). The first expression (1/2)(1+*r*) is the inclusive fitness increment that a female gains from identical-by-descent alleles inherited by her offspring, as affected by the coefficient of relatedness between the female and the sire of her offspring (*r*) scaled by 1/2 to give each parent's genetic contribution to its offspring. The second expression (1 − exp [−*c*(*m* − *m*_*min*_ − *βr*)]) is the individual offspring's viability, which is affected by *m* and *r*, and by the shape of the curve relating PI to offspring viability (*c*; i.e., how 'diminishing' returns are in *ζ*_off_ with increasing *m*). To avoid having offspring with negative viability, we constrain Eq. 1 to apply only when *m* > *m*_*min*_ + *βr*, where *m*_*min*_ is a minimum value for offspring viability given outbreeding and *β* is the magnitude of ID that can potentially be mitigated by PI; where this condition does not apply, offspring viability (and therefore *ζ*_off_) equals zero. When a focal female inbreeds, the first expression increases because more identical-by-descent alleles are inherited by inbred offspring, but the second expression decreases if *β* > 0 due to ID. However, increased PI (*m*) can offset ID and thereby increase *ζ*_off_. For simplicity, multiplication of the first and second expressions assumes statistical independence between offspring viability and relatedness to the focal female.

**Table 1:**
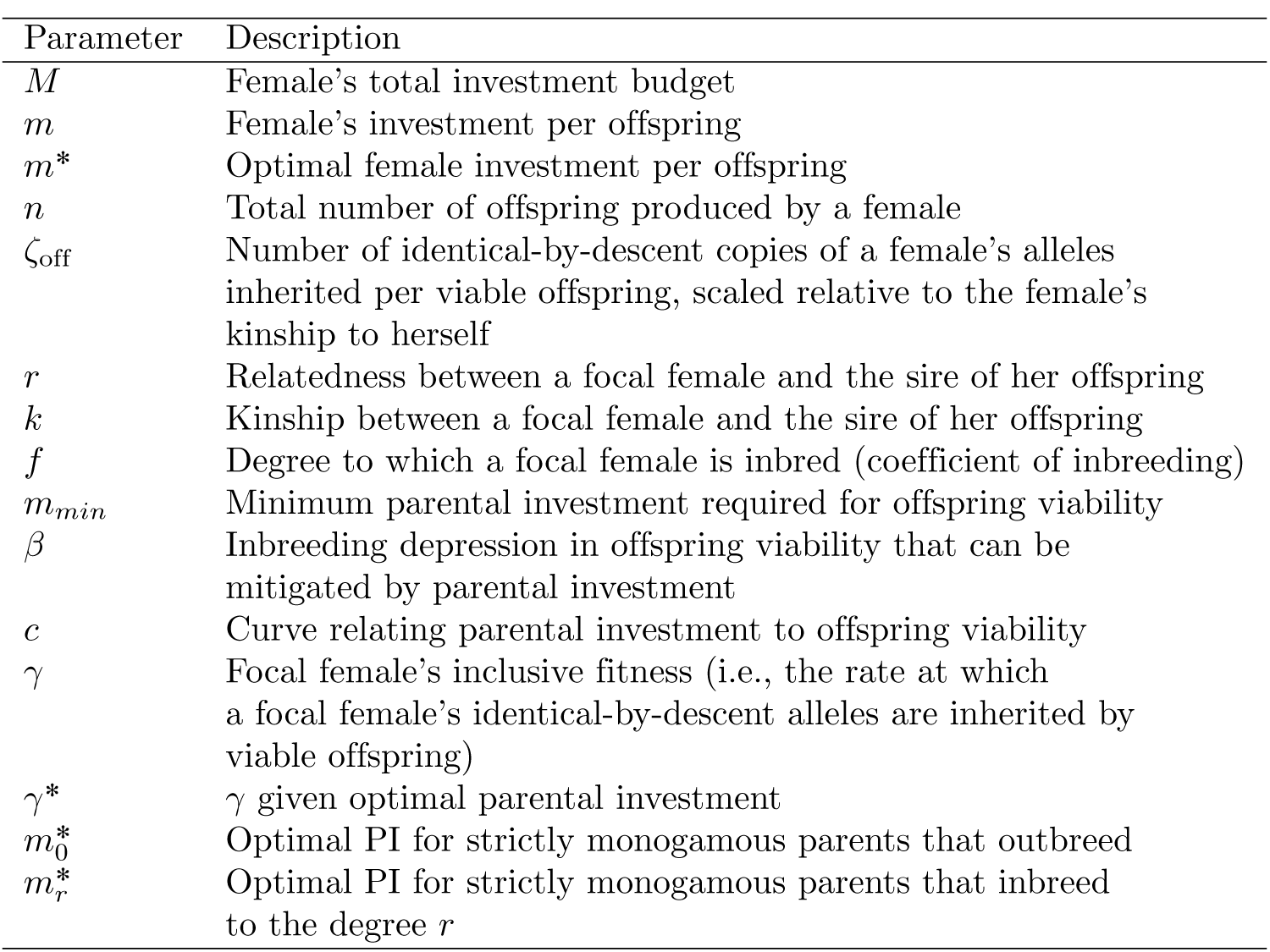
Definitions of key parameters.

When *r* = 0 and *β* = 0, Eq. 1 reduces to standard models of PI that assume outbreeding (e.g., Macnair and Parker, 1978; Parker and Macnair, 1978), but with the usual parameter *K* replaced by 1/2, thereby explicitly representing identical-by-descent alleles instead of an arbitrary constant affecting offspring fitness. Similarly, given δ = exp[−*c*(*m* − *m*_*min*_ − *βr*)], Eq. 1 reduces to standard models of biparental inbreeding that assume PI is fixed, where *δ* defines the reduced viability of inbred versus outbred offspring (see Kokko and Ots, 2006; Parker, 2006; Duthie and Reid, 2015). All offspring have equal viability as *m*→ ∞ because *β* is specifically defined as ID that can be mitigated by PI. If some additional component of ID exists that cannot be mitigated by PI, the inclusive fitness of inbreeding females decreases, but optimal PI remains unchanged (see Supporting Information p. S1-2).

## Parental investment and fitness given inbreeding

Equation 1 can be analysed to determine optimal PI (*m*\*), and a focal female's corresponding inclusive fitness *γ* given *m*\* (Kuijper and Johnstone, 2012), which we define as *γ*\*. Before analysing Eq. 1 generally, we provide a simple example contrasting outbreeding (*r* = 0) with inbreeding between first order relatives (*r* = 1/2). For simplicity, we set parameter values equal to *m*_*min*_ = 1, *β* = 1, and *c* = 1 (see Appendix 1 for example derivations of *m*\* and *γ*\* under these conditions).

Figure 1A shows how *ζ*_off_ increases with *m* given *r* = 0 (solid curve) and *r* = 1/2 (dashed curve). Increasing *r* increases the minimum amount of PI required to produce a viable offspring. Nevertheless, because inbred offspring inherit more identical-by-descent copies of their parent's alleles, sufficiently high *m* causes *ζ*_off_ of inbred offspring to exceed that of outbred offspring. The point on the line running through the origin that is tangent to *ζ*_off_(*m*) defines optimal PI (*m*\*). Figure 1A shows that for outbreeding 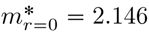 (solid line), whereas for inbreeding with a first order relative 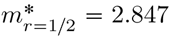 (dashed line). The slope of each tangent line identifies *γ* given optimal PI under outbreeding 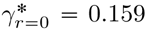 and first order inbreeding 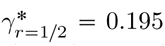. To maximise their inclusive fitness, females that inbreed with first order relatives should therefore invest more in each offspring than females that outbreed 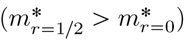. This result is general across different values of *r*; as *r* increases, so does *m*\* (see Appendix 2). Given the trade-off between *m* and *n*, females that inbreed more should therefore invest more per capita in fewer total offspring.

**Figure 1:**
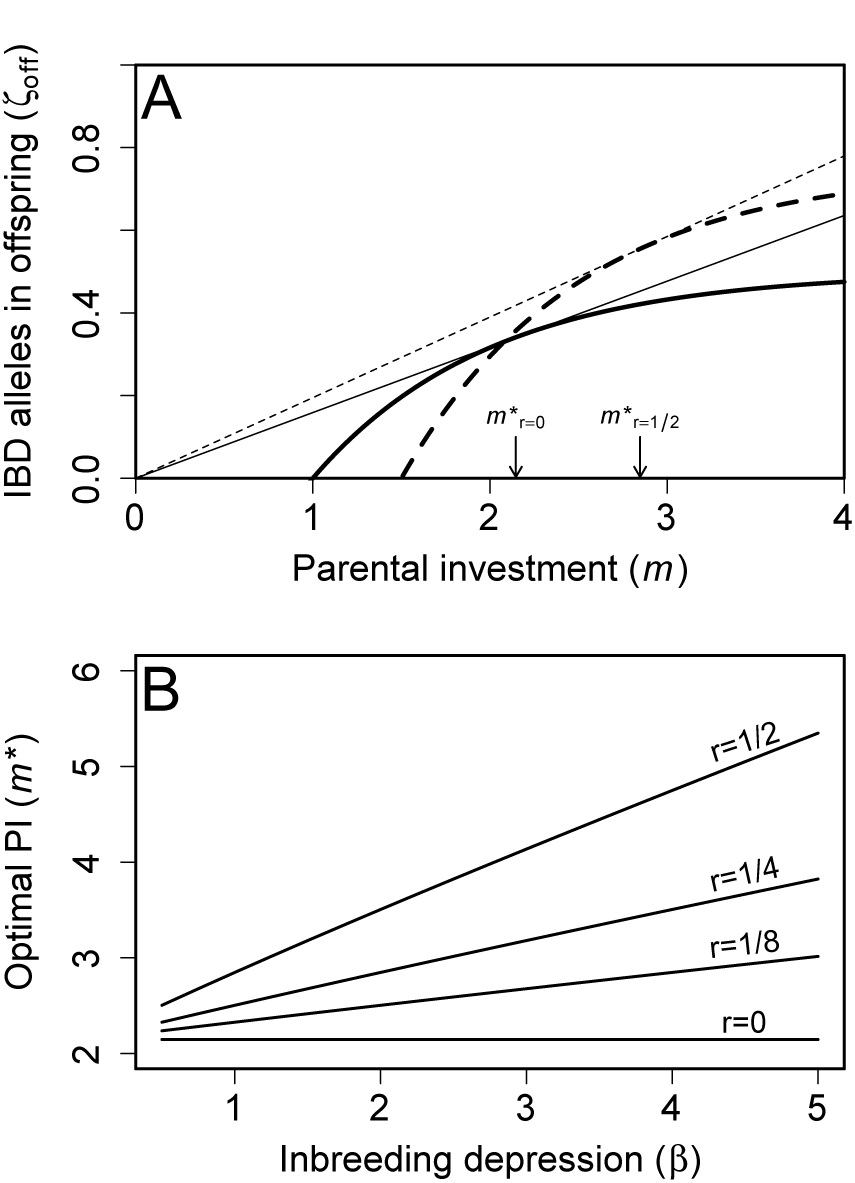
(A) Relationship between parental investment per offspring (*m*) and the number of identical-by-descent (IBD) copies of a focal female's alleles that are inherited by its offspring (*ζ*_off_) for females that outbreed (*r* = 0; solid curve) and females that inbreed with a first order relative (*r* = 1/2; dashed curve). Tangent lines identify optimal parental investment, and their slopes define a female's inclusive fitness when outbreeding (solid line) and inbreeding with a first order relative (dashed line). (B) Relationship between the magnitude of inbreeding depression in offspring viability (*β*) and optimal parental investment (*m*\*) across four degrees of relatedness (*r*) between a focal female and the sire of her offspring. Across all *β* and *r* presented, *m*_*min*_ = 1 and *c* = 1.

A general relationship between *β* and *m*\* for different values of *r* can be determined numerically. Figure 1B shows how *m*\* increases with increasing *β* and *r*, and shows that the difference in optimal PI per offspring is often expected to be high for females that inbreed with first order relatives (*r* = 1/2) rather than outbreed (e.g., when *β* = 3.25, optimal PI doubles, 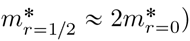.

Assuming that females allocate PI optimally, their *γ*\* values can be compared across different values of *r* and *β*. For example, given *r* = 0 and *r* = 1/2 when *β* = 1, females that inbreed by *r* = 1/2 increase their inclusive fitness more than females that outbreed (*r* = 0) when both invest optimally 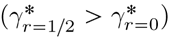. This result concurs with biparental inbreeding models where PI does not vary (see Supporting Information p. S1-4). However, if *β* = 3, then 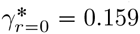 and 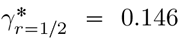. Given this higher *β*, females that outbreed will therefore have higher inclusive fitness than females that inbreed with first order relatives. Figure 2A shows more generally how *γ*\* changes with *β* and *r* given optimal PI. Across all *β*, the highest *γ*\* occurs either when *r* = 1/2 (*β* < 2.335) or *r* = 0 (*β* > 2.335), and never for intermediate values of *r*. If females can invest optimally, it is therefore beneficial to either maximise or minimise inbreeding, depending on the strength of ID.

In some populations, individuals might be unable to discriminate between relatives and non-relatives, and hence unable to facultatively adjust PI when inbreeding. We therefore consider a focal female's inclusive fitness when she cannot adjust her PI, and *m* is instead fixed at optimal PI given outbreeding. Figure 2B shows that when inbreeding females allocate PI as if they are outbreeding, *γ* always decreases, and this inclusive fitness decrease becomes more severe with increasing *r*. While the inclusive fitness of an optimally investing female that inbreeds with a first order relative (*r* = 1/2) exceeds that of an outbreeding females when *β* < 2.335, if the inbreeding female invests at the outbreeding female's optimum, then her inclusive fitness is higher only when *β* < 1.079. Consequently, if parents are unable to recognise that they are inbreeding and adjust their PI accordingly, their inclusive fitness might be decreased severely relative to optimally investing parents.

**Figure 2:**
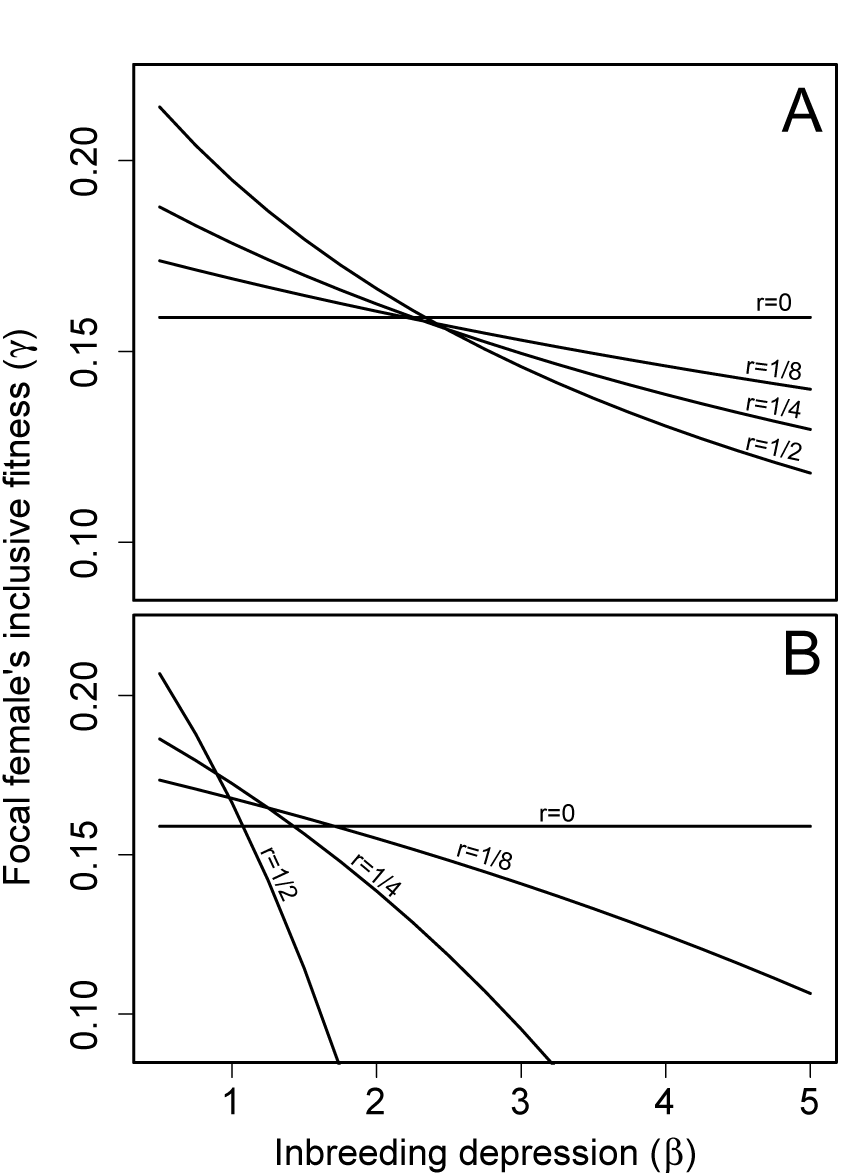
Relationship between the magnitude of inbreeding depression (*β*) and a focal female's inclusive fitness (*γ*) across four degrees of relatedness (*r*) between a focal female and the sire of her offspring assuming that focal females (A) invest optimally given the degree to which they inbreed and (B) invest at the optimum for outbreeding.

## Investment and fitness of an inbred female

Our initial assumption that a focal female is herself outbred is likely to be violated in populations where inbreeding is expected to occur (Duthie and Reid, 2015). We therefore consider how the degree to which a focal female is herself inbred will affect her optimal PI (*m*\*) and corresponding inclusive fitness (*γ*\*).

To account for an inbred female, we decompose the coefficient of relatedness *r* into the underlying coefficient of kinship *k* between the female and the sire of her offspring and the female's own coefficient of inbreeding *f* (see Hamilton, 1972; Michod and Anderson, 1979), such that,

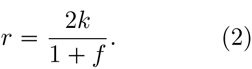

The coefficient *k* is the probability that two homologous alleles randomly sampled from the focal female and the sire of her offspring are identical-by-descent, while *f* is the probability that two homologous alleles within the focal female herself are identical-by-descent. The value of *k* between two parents therefore defines offspring *f*. Because ID is widely assumed to be caused by the expression of homozygous deleterious recessive alleles and reduced expression of overdominance (Charlesworth and Willis, 2009), *k* determines the degree to which ID is expressed in offspring. In contrast, *f* does not directly affect the degree to which homologous alleles will be identical-by-descent in offspring, and therefore does not contribute to ID. To understand how *ζ*_off_ is affected by *f* and *k*, and thereby relax the assumption that a focal female is outbred, we expand Eq. 1,

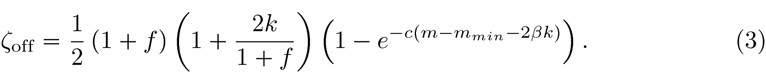

Because a focal female's *f* does not affect ID in its offspring, and instead only affects the inclusive fitness increment 1/2(1+*f*)(1+2*k*/[1+*f*]), *m*\* is unaffected by *f* (see Appendix 2). The degree to which a female is herself inbred therefore does not affect optimal PI (Fig. 3). It is worth noting that this prediction does not assume that inbred and outbred females have identical total investment budgets (*M*). Because *M* does not appear in the calculation of *m*\* (Appendix 1), an inbred focal female will have the same optimal PI as an outbred focal female even if she has a lower *M* and consequently produces fewer offspring (*n*).

Further, a focal female's *f* should only slightly affect *γ*\*. Figure 3 shows that when *k* = 1/4, *γ*\* is slightly higher for inbred females whose parents were first-order relatives (*f* = 1/4; dot-dash curve) than it is for outbred females (*f* = 0; solid curve).

**Figure 3:**
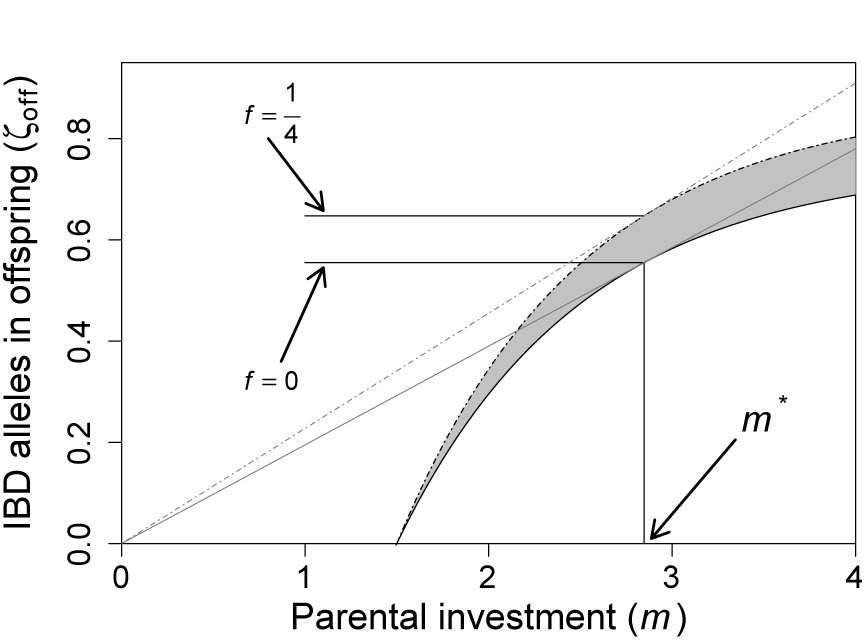
Relationship between parental investment (*m*) and the number of identical-by-descent (IBD) copies of a focal female's alleles that are present in its viable offspring (*ζ*_off_) for females that are outbred (*f* = 0; solid lower curve) versus females that are inbred (*f* = 1/4; dot-dashed upper curve). Grey shading between the curves shows the difference in *ζ*_off_ between outbred and inbred females across different degrees of parental investment. Thin grey tangent lines for each curve identify optimal parental investment (*m*\*).

## Effects of biparental investment

Our initial model assumed that only females provide PI. We now consider the opposite extreme, where PI is provided by two parents that pair exactly once in life and therefore have completely overlapping fitness interests (i.e., strict monogamy; Parker, 1985). Given Parker's (1985) implicit assumption of outbreeding, optimal PI per parent (*m*\*) does not differ between female-only PI versus monogamy (i.e., biparental investment), but twice as many offspring are produced due to the doubled total investment budget 2*M*. However, *m*\* given monogamy will differ from *m*\* given female-only PI if monogamous parents are related. This is because a male is by definition precluded from mating with another female, and therefore pays a complete opportunity cost for inbreeding (Waser et al., 1986). A focal female will thereby lose any inclusive fitness increment that she would have otherwise received when her related mate also bred with other females.

To incorporate this cost, we explicitly consider both the direct and indirect fitness consequences of inbreeding. We assume that if a focal female avoids inbreeding with her male relative, then that relative will outbreed instead, and that parents are outbred (*f* = 0) and invest optimally for any given *β*. We define 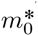 and 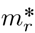 as optimal investment for outbreeding and for inbreeding to the degree *r*, respectively. Therefore, if a focal female avoids inbreeding,

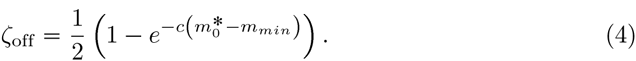

If she instead inbreeds,

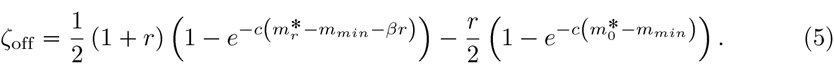

The first term of Eq. 5 represents the fitness increment the focal female receives from inbreeding. The second term represents the indirect loss of fitness the focal female would have received through her male relative had she not inbred with him. The resulting decrease in *ζ*_off_(*m*_*r*_) causes an overall increase in 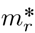. All else being equal, monogamous parents should therefore each invest even more per offspring when inbreeding than females should invest given female-only PI. For example, if *r* = 1/2 and *β* = 1, 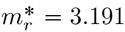 given strict monogamy but 2.847 given female-only PI (Fig. 4A). However, if *r* = 1/2, then 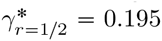 given female-only PI, but 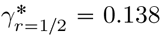 given strict monogamy. The latter is therefore less than the increase resulting from optimal PI given outbreeding, 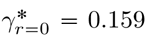. Indeed, given strict monogamy, 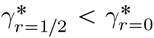, meaning that inclusive fitness accrued from inbreeding never exceeds that accrued from outbreeding. While these calculations assume that a male relative would otherwise outbreed, Duthie and Reid (2015) illustrate how this assumption might be relaxed.

**Figure 4:**
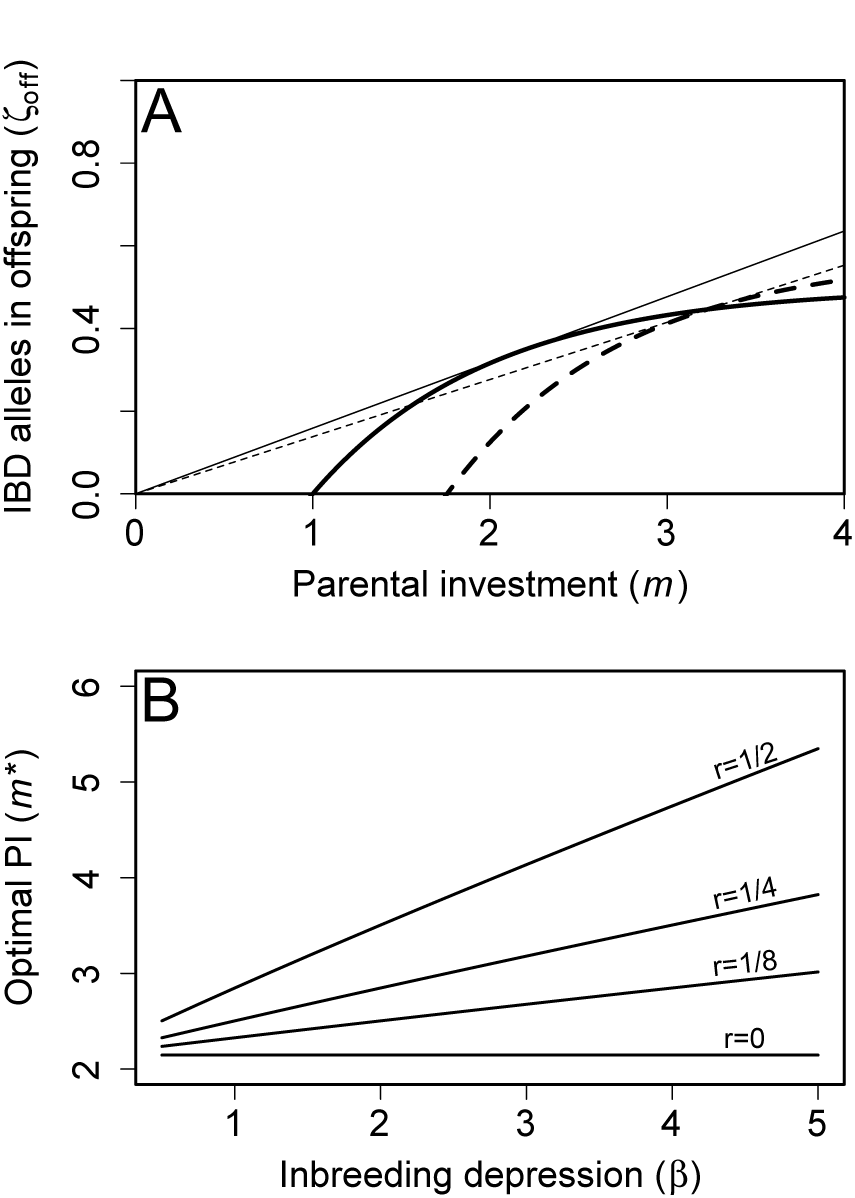
Assuming strict monogamy, the (A) relationship between parental investment (*m*) and the number of identical-by-descent (IBD) copies of a focal female's alleles that are inherited by its viable offspring (*ζoff*) for females that outbreed (*r* = 0; solid curve) and females that inbreed with first order relatives (*r* = 1/2; dashed curve). Tangent lines identify optimal parental investment (*m*\*), and their slopes (*γ*\*) define a female's inclusive fitness when outbreeding (solid line) and inbreeding (dashed line). (B) Relationship between the magnitude of inbreeding depression (*β*) and *m*\* across four degrees of relatedness (*r*) between a focal female and the sire of her offspring.

Figure 4A shows how *ζ*_off_ increases as a function of *m* given *r* = 0 (solid curve) and *r* = 1/2 (dashed curve) given strict monogamy, and can be compared to analogous relationships for female-only PI, given identical parameter values shown in Fig. 1A. In contrast to female-only PI, 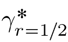 (slope of the dashed line) is now lower when *r* = 1/2 than when *r* = 0, meaning that the inclusive fitness of females that inbreed with first order relatives is lower than females that outbreed given strict monogamy. Figure 4B shows *m*\* for two strictly monogamous parents across different values of *r* and *β*. In comparison with female-only PI (Fig. 1B), *m*\* is always slightly higher given strict monogamy if *r* > 0 (Fig. 4B), but in both cases *m*\* increases with increasing *r* and *β*.

We now consider the inclusive fitness of focal monogamous parents that cannot facultatively adjust PI upon inbreeding, and instead assume PI is fixed at the optimum for outbreeding. Figure 5 shows how *γ* varies with *β* given that monogamous parents invest optimally and invest at an optimum PI for outbreeding. In contrast to female-only PI (Fig. 2A), *γ*\* is always maximised at *r* = 0, meaning that inbreeding never increases inclusive fitness. Inclusive fitness decreases even further when inbreeding individuals allocate PI at *m*\* for outbreeding (compare Figs. 2B and 5B). Universally decreasing *γ* with increasing *r* is consistent with biparental inbreeding theory, which demonstrates that if inbreeding with a female completely precludes a male from outbreeding, inbreeding will never be beneficial (Waser et al., 1986; Duthie and Reid, 2015). However, if relatives become paired under strict monogamy, each should invest more per offspring than given female-only PI.

**Figure 5:**
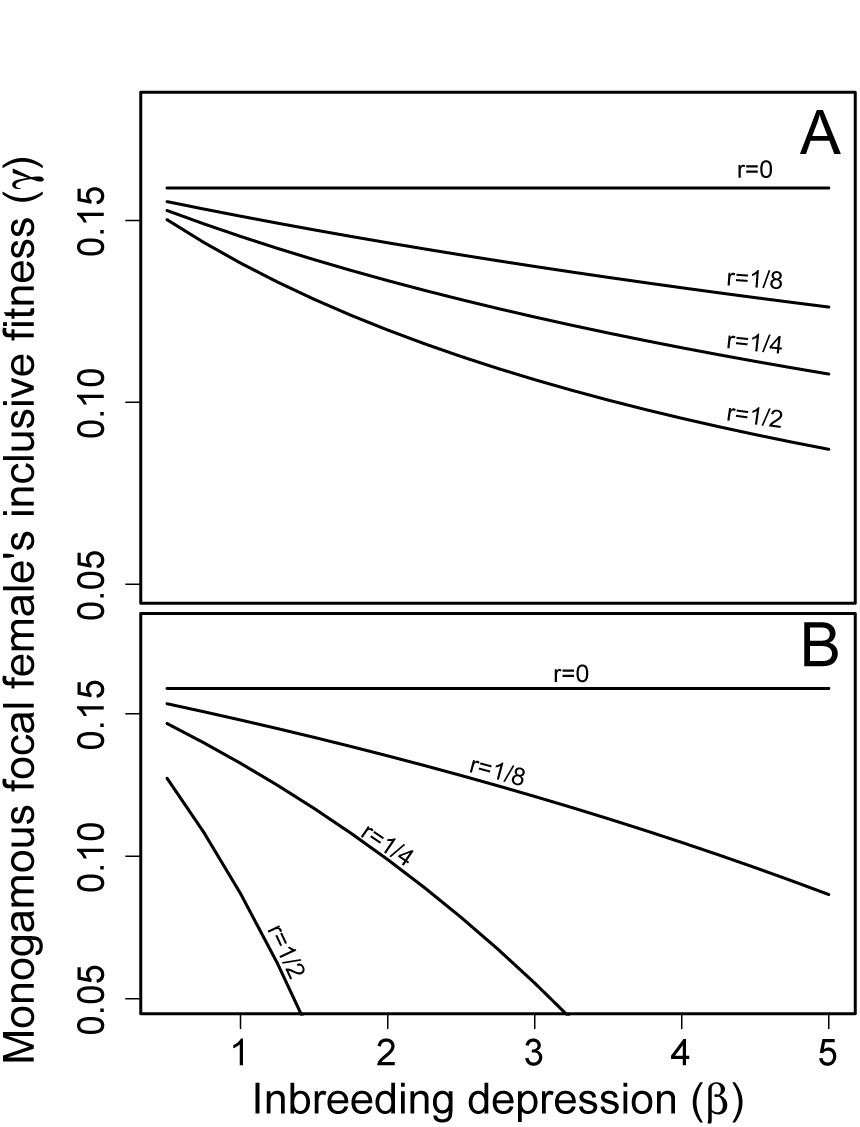
Assuming strict monogamy, the relationship between the magnitude of inbreeding depression (*β*) and a focal female's inclusive fitness (*γ*) across four degrees of relatedness (*r*) between a focal female and the sire of her offspring given that (A) focal females invest optimally given their degree of inbreeding, and (B) females invest at the optimum for outbreeding.

## Discussion

By unifying biparental inbreeding theory and parental investment (PI) theory under an inclusive fitness framework, we show that when females inbreed and hence produce inbred offspring, optimal PI always increases, and this increase is greatest when inbreeding depression in offspring viability (ID) is strong. We also show that optimal PI does not change when a focal female is herself inbred. Finally, we show that, in contrast to existing theory that implicitly assumes outbreeding (Parker, 1985), the occurrence of inbreeding means that optimal PI increases given strict monogamy and associated biparental investment compared to female-only PI. Our conceptual synthesis illustrates how previously separate theory developed for biparental inbreeding (Parker, 1979, 2006) and PI (Macnair and Parker, 1978; Parker and Macnair, 1978) can be understood as special cases within a broader inclusive fitness framework in which inbreeding and PI

## Inbreeding and PI in empirical systems

Theory can inform empirical hypothesis testing by logically connecting useful assumptions to novel empirical predictions. We demonstrate that given a small number of assumptions regarding PI and inbreeding, selection will cause PI to increase with increasing relatedness between parents and increasing magnitude of ID (Fig. 1B). The total number of offspring that inbreeding parents produce will correspondingly decrease given the fundamental trade-off with investment per offspring. Empirical studies are now needed to test key assumptions and predictions.

One key assumption is that inbreeding depression in offspring viability can be mitigated by PI. Numerous studies have estimated magnitudes of ID in components of off-spring fitness (Keller and Waller, 2002; Charlesworth and Willis, 2009; Szulkin et al., 2013). However, PI is notoriously difficult to measure because it might encompass numerous phenotypes, each affecting allocation from an unknown total PI budget (Parker et al., 2002). It is therefore difficult to quantify to what degree ID is reduced by PI, and few empirical studies have estimated such effects. One technique might be to experimentally remove parents during offspring development, thereby precluding any PI expressed through parental care (note that ‘PI’ is not synonymous with ‘parental care’; the latter refers to any parental phenotype that increases offspring fitness, and is not necessarily defined by a trade-off with number of offspring, Gardner and Smiseth, 2011; Royle et al., 2012). For example, Pilakouta et al. (2015) quantified the fitness of inbred and outbred burying beetle (*Nicrophorus vespilloides*) offspring in the presence and absence of maternal care, finding that maternal care increased survival of inbred offspring relatively more than survival of outbred offspring (see also Pilakouta and Smiseth, 2016). Interpreting care as a component of PI, this result concurs with the assumption that PI can reduce ID. Similarly, in the subsocial spider *Anelosimus* cf. *jucundus*, in which care is provided by solitary females, Avilés and Bukowski (2006) found evidence of ID only late in an offspring's life when parental care was no longer provided, and hypothesised that care might buffer ID. However, *A.* cf. *jucundus* females that inbred did not produce fewer offspring than females that outbred, as our model predicts if females respond to inbreeding by increasing PI. Some further constraint might therefore prevent female *A.* cf. *jucundus* from adaptively adjusting PI.

Indeed, a second assumption predicting optimal PI is that individuals can discriminate among different kin and non-kin and adjust their PI according to the degree to which they inbreed. While kin discrimination has been observed in multiple taxa (e.g., Mateo, 2002; Griffin and West, 2003; Dudley and File, 2007; Strassmann et al., 2011), if parents are unable to infer that they are inbreeding, they will likely allocate PI sub-optimally, resulting in decreased fitness of inbreeding parents (Fig. 2B) and decreased viability of resulting inbred offspring. The realised magnitude of ID might consequently be greater than if PI were allocated optimally, implying that observed ID depends partly on adaptive PI rather than resulting solely from offspring homozygosity and inbreeding load. To our knowledge, no empirical studies have explicitly tested whether or not PI varies with inbreeding. However, strong negative correlations between the degree to which parents inbreed and litter size have been found in wolves (*Canis lupus*; Liberg et al., 2005; Fredrickson et al., 2007). Wolves are highly social and generally monogamous, and are likely able to discriminate among kin (Räikkönen et al., 2009; Geffen et al., 2011). Liberg et al. (2005) and Fredrickson et al. (2007) interpret decreased litter size as a negative fitness consequence of inbreeding manifested as increased early mortality of inbred offspring. Our model suggests an alternative explanation; smaller litter sizes might partially reflect adaptive allocation whereby inbreeding parents invest more in fewer offspring. Future empirical assessments of the relative contributions of ID and adjusted PI in shaping offspring viability, and tests of the prediction that inbreeding parents should produce fewer offspring, will require careful observation of variation in PI and litter or brood sizes in systems with natural or experimental variation in inbreeding.

Our model also clarifies why an individual's reproductive success, simply measured as the number of offspring produced, does not necessarily reflect inclusive fitness given inbreeding, or hence predict evolutionary dynamics. A female that produces an outbred brood might have lower inclusive fitness than a female that produces an inbred brood of the same (or slightly smaller) size if the inbreeding female's viable offspring carry more identical-by-descent allele copies (see also Reid et al., 2016). Interestingly, if brood size is restricted by some physiological or external constraint (i.e., brooding or nest site capacity), our model predicts that females with large total resource budgets *M* might benefit by inbreeding and thereby adaptively allocate more PI to each offspring. Overall, therefore, our model shows that understanding the evolutionary dynamics of reproductive systems that involve interactions among relatives is likely to require ID, inbreeding strategy, and reproductive output to be evaluated in the context of variable PI.

## Intrafamilial conflict given inbreeding

Interactions over PI are characterised by intrafamilial conflict between parents, between parents and offspring, and among siblings (Parker et al., 2002). Our general theoretical framework sets up future considerations of intrafamilial conflict over PI given inbreeding. Our current model assumes either female-only PI or strict monogamy, the latter meaning that female and male fitness interests are identical, eliminating sexual conflict. However, in general, if both parents invest and are not completely monogamous, sexual conflict is predicted because each parent will increase its fitness if it provides less PI than its mate. Optimal PI can then be modelled as an evolutionary stable strategy (Maynard Smith, 1977), and is expected to decrease for both parents (Parker, 1985). This decrease in optimal PI might be smaller given inbreeding because the negative inclusive fitness consequences of a focal parent reducing PI will be exacerbated if the mate that it abandons is a relative.

Sexual conflict might also be minimised if a focal parent that decreases its PI must wait for another mate to become available before it can mate again. Kokko and Ots (2006) considered the fitness consequences of inbreeding and inbreeding avoidance given a waiting time between mate encounters, and a processing time following mating, which they interpreted as PI. They found that inbreeding tolerance generally increased with increasing waiting time, but that such relationships depended on processing time. However, processing time was a fixed parameter, meaning that parents could not adjust PI as a consequence of inbreeding. If this assumption was relaxed such that PI could vary, parents that inbreed might be expected to increase their time spent processing offspring before attempting to mate again, altering selection on inbreeding strategy.

Parent-offspring conflict is a focal theoretical interest of many PI models, which generally predict that offspring benefit from PI that exceeds parental optima (Macnair and Parker, 1978; Parker and Macnair, 1978; Parker, 1985; De Jong et al., 2005). However, such conflict might be decreased by inbreeding because inbreeding parents are more closely related to their offspring than are outbreeding parents. De Jong et al. (2005) modelled PI conflict in the context of optimal seed mass from the perspective of parent plants and their seeds given varying rates of self-fertilisation, predicting that conflict over seed mass decreases with increasing self-fertilisation, assuming seed mass is controlled by seeds rather than parent plants. In general, the same principles of parent-offspring conflict are expected to apply for biparental inbreeding; parent-offspring conflict should decrease with increasing inbreeding, and reduced conflict might in turn affect offspring behaviour. For example, Mattey (2014) observed both increased parental care and decreased offspring begging in an experimental study of *N. vespilloides* when offspring were inbred. A reduction in begging behaviour might be consistent with our model if it reflects decreased parent-offspring conflict due to the fitness interests of parents and inbred offspring being more closely aligned. Future models could relax our assumption that parents completely control PI, and thereby consider how inbreeding and PI interact given parent-offspring conflict.

Inbreeding increases relatedness among offspring, potentially affecting sibling conflict within or among broods (Parker et al., 2002; Bonisoli-Alquati et al., 2011; Ruch et al., 2014). Within broods, conflict over PI is directly proportional to relatedness among offspring following Hamilton's rule (Hamilton, 1964*a*,*b*), assuming that a focal female's total PI budget is fixed per brood. In contrast, the degree to which sibling conflict among broods is affected by relatedness depends on the mating system and, consequently, the degree to which PI is provided by each parent (e.g., female-only PI versus monogamy; Lessells and Parker, 1999; Parker et al., 2002). Nevertheless, because inbreeding can increase sibling relatedness both within and among broods, sibling conflict might be reduced in both cases, potentially promoting evolution of alloparental care (Gardner, 2014; Davies et al., 2016). However, the increase in PI that we predict as a consequence of inbreeding could also feed back to indirectly increase evolution of sibling competition if it increases total resources available to offspring that can be obtained from parents instead of other sources (Gardner and Smiseth, 2011).

In conclusion, we have demonstrated an intrinsic link between the effects of inbreeding and PI on inclusive fitness, thereby conceptually uniting two long-standing theoretical frameworks (Parker, 1979, 2006; Macnair and Parker, 1978; Parker and Macnair, 1978). All else being equal, parents that inbreed should produce fewer offspring so that they can invest more resources in each. Future theory might usefully incorporate our results into models of intrafamilial conflict.

## Acknowledgments

This work was funded by a European Research Council Starting Grant to JMR. We thank Greta Bocedi, Andy Gardner, Ryan Germain, Matthew Wolak, and two anonymous reviewers for helpful comments.

## Appendix 1 Example derivations of *m*\* and *γ*\*

In general, the equation for a line tangent to some function *f* at the point *a* is,

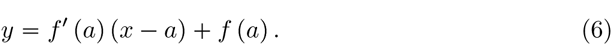

In the above, *f*′(*a*) is the first derivative of *f*(*a*), and *y* and *x* define the point of interest through which the straight line will pass that is also tangent to *f*(*a*). The original function that defines *ζ*_off_ is as follows,

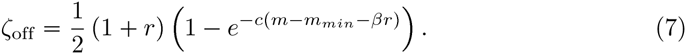

Differentiating *ζ*_off_ with respect to *m*, we have the following,

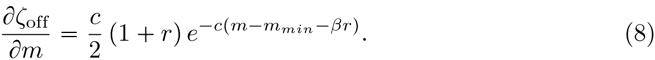

Substituting *ζ*_off_(*m*) and ∂*ζ*_off_/∂*m* and setting *y* = 0 and *x* = 0 (origin), we have the general equation,

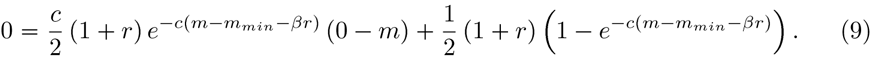

A solution for *m*\* can be obtained numerically for the example in which *m*_*min*_ = 1, *β* = 1, and *c* = 1. If *r* = 0, 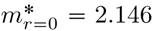, and if 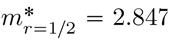. Solutions for the slopes defining 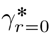 and 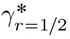 can be obtained by finding the straight line that runs through the two points (0,0) and (*m*\*, *ζ*_off_(*m*\*)). In the case of *r* = 0, *ζ*_off_(*m*\*) = 0.341, so we find, 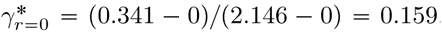. In the case of *r* = 1/2, *ζ*_off_(*m*\*) = 0.555, so we find, 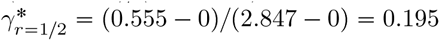.

## Appendix 2 *m*\* increases with increasing *r*

Here we show that optimal parental investment (*m*\*) always increases with increasing inbreeding given ID and *c* > 0. First, note that *m*\* is defined as the value of *m* that maximises the rate of increase in *ζ*_off_ for a female. This is described by the line that passes through the origin and lies tangent to *ζ*_off_(*m*). As in Appendix 1, we have the general equation for which *m* = *m*\*,

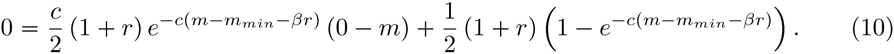

We first substitute *m* = *m*\* and note that this equation reduces to,

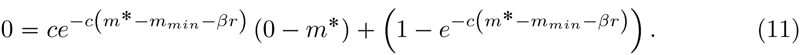

This simplification dividing both sides of the equation by (1/2)(1+*r*) has a biological interpretation that is relevant to PI. Optimal PI does not depend directly on the uniform increase in *ζ*_off_ caused by *r* in (1/2)(1+*r*), the change in *m*\* is only affected by *r* insofar as *r* affects offspring viability directly through ID.

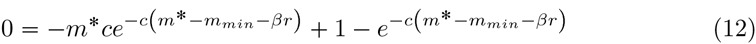

From the above, *r* can be isolated,

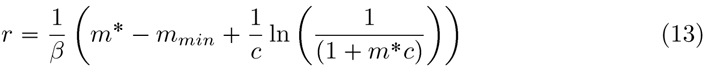

We now differentiate *r* with respect to *m*\*,

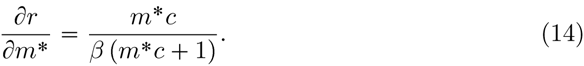

By applying the chain rule, we can thereby arrive at the general conclusion,

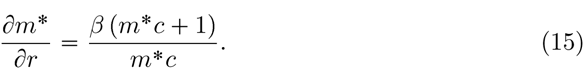

Given the above, ∂*m*\*/∂*r* > 0 assuming *β* > 0 (ID), *c* > 0 (offspring viability increases with PI), and *m*\* > 0 (optimum PI is positive). These assumptions are biologically realistic; we therefore conclude that the positive association between optimal PI (*m*\*) and inbreeding (*r*) is general. As inbreeding increases, so should optimal PI in offspring.

